# Nonivasive prenatal diagnosis of single-gene disorders using droplet digital PCR

**DOI:** 10.1101/179804

**Authors:** Joan Camunas-Soler, Hojae Lee, Louanne Hudgins, Susan R. Hintz, Yair J. Blumenfeld, Yasser Y. El-Sayed, Stephen R. Quake

**Affiliations:** Department of Bioengineering, Stanford University, Stanford, CA 94305, USA.; Division of Medical Genetics, Department of Pediatrics, Stanford University, Stanford, CA 94305, USA.; Division of Neonatal and Developmental Medicine, Department of Pediatrics, Stanford University School of Medicine, Stanford, CA 94305, USA.; Division of Maternal-Fetal Medicine and Obstetrics, Department of Obstetrics and Gynecology, Stanford University, Stanford, CA 94305, USA.; Department of Applied Physics, Stanford University, Stanford, CA 94305, USA.; Chan Zuckerberg Biohub, San Francisco, CA 94518, USA.

**Keywords:** prenatal diagnosis, cell-free DNA, single-gene disorders, biochemical disorders, noninvasive prenatal testing, liquid biopsy

## Abstract

**Background:** Prenatal diagnosis in pregnancies at risk of single-gene disorders is currently performed using invasive methods such as chorionic villus sampling and amniocentesis. This is in contrast with screening for common aneuploidies, for which noninvasive methods with a single maternal blood sample have become standard clinical practice.

**Methods:** We developed a protocol for noninvasive prenatal diagnosis of inherited single gene disorders using droplet digital PCR (ddPCR) from circulating cell-free DNA (cfDNA) in maternal plasma. First, the amount of cfDNA and fetal fraction are determined using a panel of Taqman assays targeting high-variability SNPs. Second, the ratio of healthy and diseased alleles in maternal plasma are quantified using Taqman assays targeting the mutations carried by the parents. Two validation approaches of the mutation assay are presented.

**Results:** We collected blood samples from 9 pregnancies at risk for different single gene disorders including common conditions and rare metabolic disorders. We measured cases at risk of hemophilia, ornithine transcarbamylase deficiency, cystic fibrosis, β-thalassemia, mevalonate kinase deficiency, acetylcholine receptor deficiency and DFNB1 nonsyndromic hearing loss. We correctly differentiated affected and unaffected pregnancies (2 affected, 7 unaffected), confirmed by neonatal testing. We successfully measured an affected pregnancy as early as week 11, and with a fetal fraction as low as 3.7±0.3%.

**Conclusion:** Our method detects single nucleotide mutations of autosomal recessive diseases as early as the first trimester of pregnancy. This is of importance for metabolic disorders where early diagnosis can affect management of the disease and reduce complications and anxiety related to invasive testing.

## Introduction

The presence of circulating cell-free DNA (cfDNA) of fetal and placental origin in maternal plasma has allowed the development of noninvasive tools to detect fetal genetic abnormalities from a maternal blood draw ^1–3^. Currently, noninvasive prenatal testing (NIPT) of common aneuploidies (e.g. Down syndrome) is clinically available as a screening test that can be performed as early as week 10 of pregnancy with false positive rates below 0.2% ^2,4,5^ and without the complications related to invasive testing ^6,7^. More recently, NIPT has also become commercially available for some genomic microdeletions (e.g. DiGeorge syndrome, Cri-du-chat syndrome) ^8–10^.

However, prenatal diagnosis of pregnancies at risk of single gene disorders still requires the use of invasive techniques such as amniocentesis or chorionic villus sampling (CVS). These methods have a risk of miscarriage, cause higher discomfort, and can only be applied during certain time windows of pregnancy ^6,7^. Although commercial development of screening tests for single gene disorders is difficult due to the low prevalence of each given mutation in the general population (hampering positive predictive values), the development of accurate NIPT could replace invasive testing and become a diagnostic test for parents who are carriers of a mutation, who are at high risk of having an affected pregnancy ^11^. A recent carrier screening study including >350,000 individuals has shown that the number of pregnancies at risk of 94 common severe single gene disorders ranges between 95 to 392 per 100,000 depending on ethnic background ^12^. Remarkably, these values are comparable to the prevalence of common aneuploidies such as trisomy 21 ^13^.

The development of noninvasive tools for these disorders is important as it allows patients and doctors to make informed decisions in pregnancies at risk of severe conditions while reducing anxiety related to invasive or postnatal testing. In addition early treatment is sometimes available for conditions that might otherwise cause irreversible damage to the fetus such as metabolic disorders or congenital malformations (e.g. dietary treatment or neonatal surgery respectively) ^14–16^. Finally, prenatal diagnostics might also prove useful to develop protocols for cord blood collection in views of potential cures of inherited single-gene disorders by using gene-editing techniques on hematopoietic stem cells ^17,18^.

Detection of single-gene disorders is straight-forward for paternally-inherited mutations or common de novo mutations, where the presence of a mutated allele in maternal plasma can be directly attributed to an affected fetus and not to background cfDNA of maternal origin ^19–22^. However, most common single-gene disorders are autosomal-recessive due to their deleterious nature, and therefore one must carefully quantitate the ratio of mutant to wild type alleles in order to genotype the fetus. This problem has been solved in principle by applying the counting principle to high depth whole exome sequence or to full haplotypes ^23^, but this approach requires the use of sequencing and is more costly than digital PCR. Proof-of-concept studies with digital PCR have been conducted for a number of autosomal recessive and X-linked disorders ^24–30^, but a general method to perform noninvasive diagnosis of these conditions is not yet available. Previous digital PCR studies have been limited in that they have not had large enough SNP panels to measure the fetal fraction in the general population, or have not had enough SNP measurements to estimate the error in measurement of fetal fraction.

Here we address these challenges by developing a simple droplet digital PCR (ddPCR) protocol to diagnose autosomal and X-linked single gene disorders. This protocol is applied directly to the maternal cell free DNA sample and does not require a separate maternal genotyping step. An accurate quantification of the fetal fraction is achieved by targeting a panel of 47 high-variability SNPs, and we show that the final measurement error in determining fetal genotype is composed of roughly equal contributions from the error in fetal fraction and the Poisson error due to counting statistics. We show how this method enables diagnosis of recessive single gene disorders, both when they are due to a mutation shared by both progenitors or to heterozygous compound mutations (when father and mother carry a different mutation affecting the same gene). Unambiguous results are shown for samples with a fetal fraction as low as 3.6%.

## Results

### Clinical protocol and validation of assays

In this study we enrolled pregnant patients who are carriers of mutations causing autosomal recessive or X-linked disorders. We then followed the experimental protocol depicted in Figure 1 to test whether the fetus is affected by the disease. For each pregnancy at risk of a known mutation, we designed primers to amplify the region of the mutation and TaqMan probes labeled with different fluorophores against the healthy and mutated allele at-risk (i.e. single nucleotide mutation, insertion, deletion). The assays were validated using genomic DNA (gDNA) of carriers and non-carriers of the mutation in ddPCR experiments (Supp. Section S1 and Supp. Fig. S1). Carrier gDNA was obtained from nucleated cells from maternal blood. Once we enrolled a patient carrying a new target mutation, the limiting factor to perform a diagnostic was the ordering time of the Taqman assays (∼1 week). In order to be able to validate the assays before maternal blood collection, we developed an alternative approach using mixtures of synthetic DNA fragments and spike in experiments (Fig. 2 and Supp. Section S1). This allowed us to validate the Taqman probes before sample collection and reduce the turnaround time of the assay to ∼1 day.

**Figure 1.**
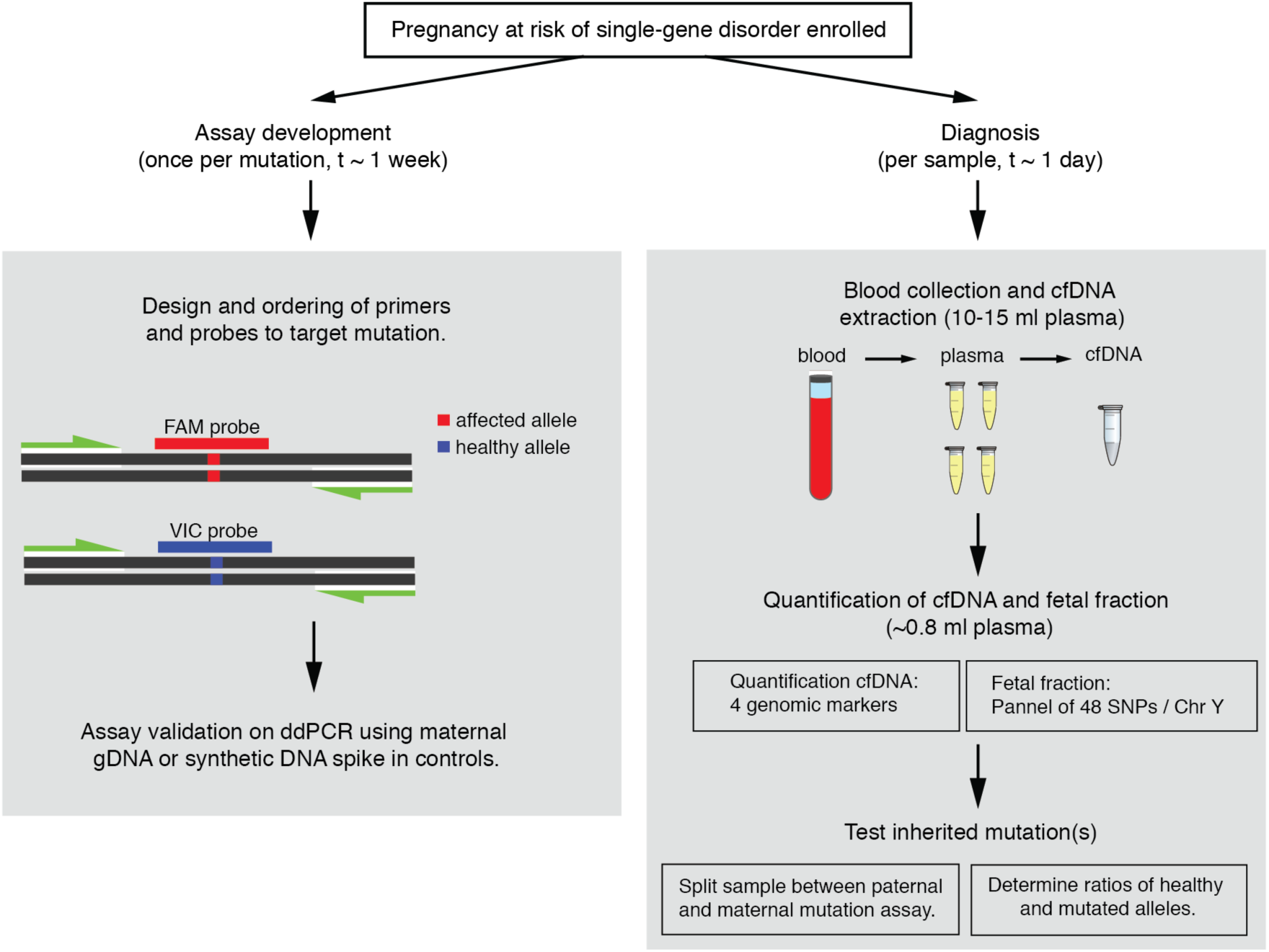
Protocol for noninvasive prenatal diagnostics of single-gene disorders.

**Figure 2.**
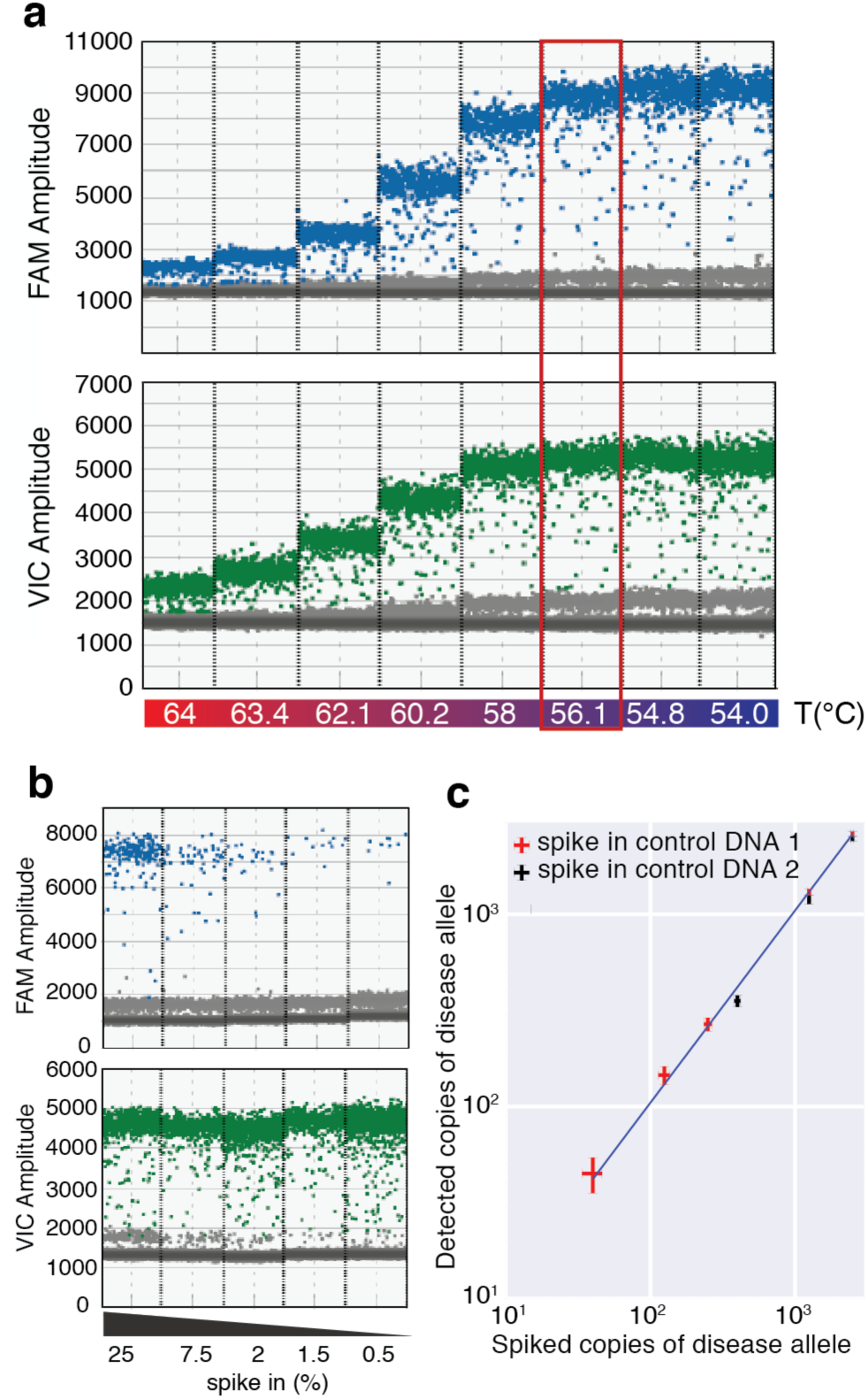
Validation of diagnostic assays with synthetic spike in controls (g-bocks). **(a)** Temperature gradient of 1:1 mixtures of synthetic DNA fragments containing the mutant (FAM) and healthy (VIC) allele for mutation c.835C>T in OTC gene (dbSNP: rs72558455). The optimal temperature for the Taqman assay in ddPCR experiments is highlighted in red. **(b)** Spike in controls of the synthetic mutant allele (FAM) in fragmented genomic DNA of a healthy donor. Scatter plots of FAM/VIC fluorescence are shown in Supp. Fig. S2. **(c)** Quantification using ddPCR of varying amounts of spike in synthetic DNA (mutant allele) in a background of fragmented gDNA (∼5000 genome equivalents per reaction) from two different healthy donors (red, black). Error bars are obtained from Poisson statistics.

For each incoming sample, we extracted cfDNA from ∼30 ml of maternal blood (see Methods and Figure 1). We then performed a quantification assay of the total amount of cfDNA and fetal fraction using Taqman assays targeting 4 genomic markers (cfDNA quantification), and a panel of 47 high-variability SNPs and a X/Y chromosome marker (fetal fraction determination, see Methods). This information was used to decide if a determinative result was possible and to determine the optimal split of sample to test the paternally and maternally-inherited mutations in compound heterozygous conditions, as well as the confidence intervals of the result.

### Quantification of cfDNA and fetal fraction

We used approximately 7% of each sample to quantify the fetal fraction and total amount of cfDNA (see Methods). For each sample we measured the minor allele fraction (MAF) for each SNP in the panel and determined the fetal fraction from the distribution of SNPs that are homozygous for the mother and heterozygous for the fetus, which are found in the range 0.5< MAF<15 (Fig. 3a and Methods). These SNPs are the most informative to determine the fetal fraction as the measured copies from the fetal allele are not affected by background noise coming from the more abundant maternal alleles. Interestingly, our assay also allowed us to discriminate SNPs that are heterozygous for the mother but homozygous for the fetus, which show a characteristic symmetric peak in the range 35< MAF< 50 (Fig. 3a). Although we did not use these SNPs to calculate the fetal fraction due to its higher noise, they could also be used to improve the estimate if a reduced SNP panel is used (Supp. Fig. S3). The total quantification of cfDNA was also obtained for each sample, as well as the sex of the fetus (Fig. 3a, insets). We compared the standard deviation of the fetal fraction measurement to the expected noise due Poisson subsampling (as we are using a limited amount of sample for this measurement), finding a good agreement between experimental measurement and theoretical expectation for all samples. This is relevant for samples with a low fetal fraction or total amount of cfDNA, where additional noise in the fetal fraction measurement would affect the confidence intervals of the diagnostic assay. As expected, the fetal fraction increases with gestational age (Fig. 3a), a result that is also consistently observed for individual SNPs of the panel (Supp. Fig. S4). Finally, results for 12 different pregnancies show a distribution of maternal and fetal genotypes compatible with the expected values for high-variability SNPs, suggesting that our panel can be used to determine the fetal fraction in populations of different genetic background (Fig. 3b and Methods).

**Figure 3.**
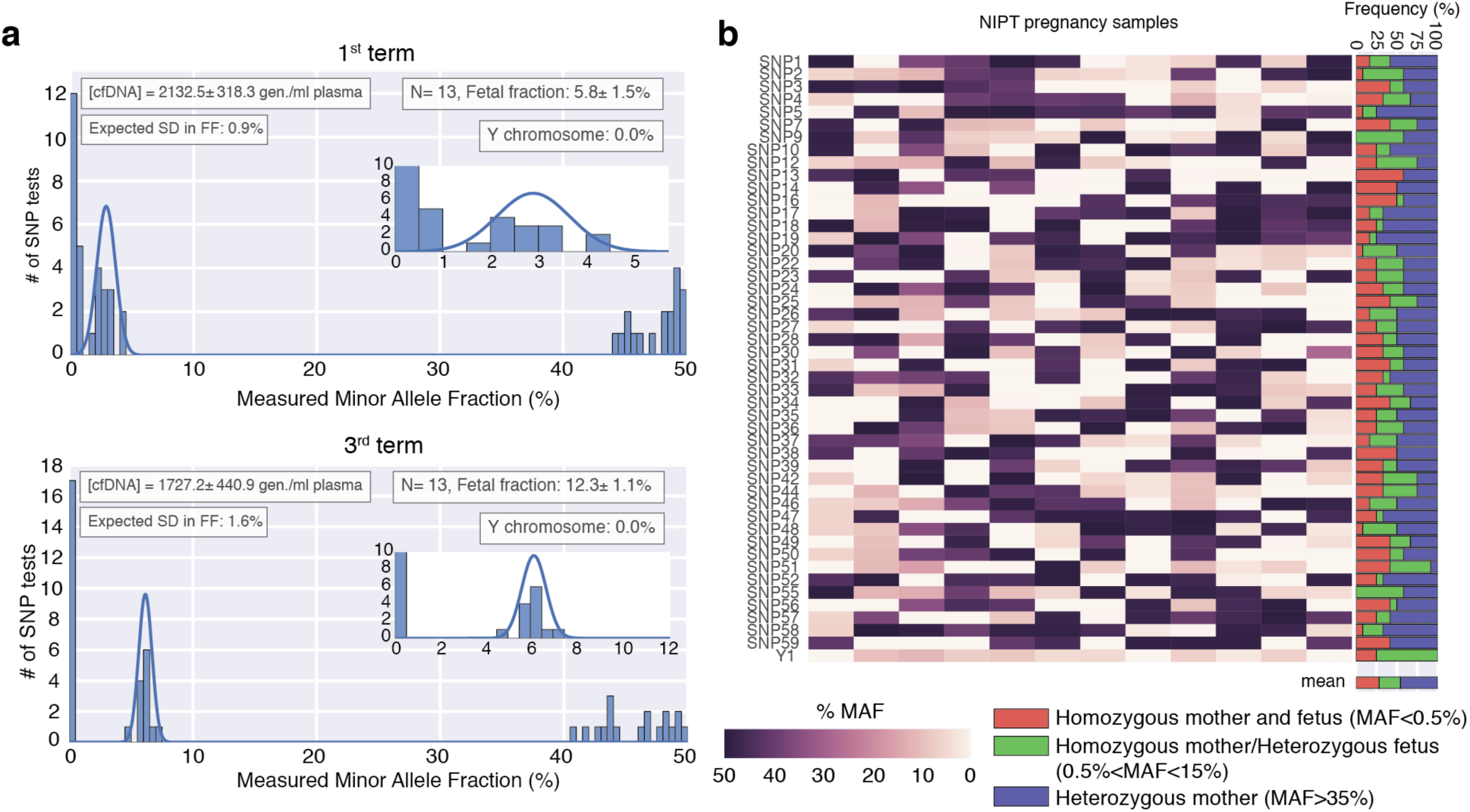
Quantification of cfDNA and fetal fraction determination. **(a)** Histogram of the MAF for the 47 SNP assays used to determine the fetal fraction. Top (bottom) panel are results from a first (third) term sample of the same pregnancy. The fetal fraction is determined from SNPs that are homozygous for the mother and heterozygous for the fetus (found in the range 0.5%< MAF<15%) and calculated as 2*MAF. A gaussian fit to these SNPs is shown in blue. Inset boxes show the (i) quantification of cfDNA in the sample (determined as explained in the Methods section) (ii) the fetal fraction and number of informative SNP assays (*N*), (iii) expected error in the fetal fraction as explained in the methods section, and (iv) sex determination assay. Errors are reported as standard deviation. **(b)** MAF of the 47-SNP assay for 12 different pregnancies. The right panel shows the frequency of each combination of maternal and fetal genotypes. The recovered distributions are in agreement with the expected results for high-variability SNPs (Heterozygous mother ∼50%, Homozygous mother and fetus ∼25%, Homozygous mother/Heterozygous fetus ∼25%).

### Diagnosis of X-linked disorders and autosomal recessive disorders

We first addressed the case of X-linked mutations, where the carrier status of the mother poses a risk for pregnancies carrying a male fetus. We analyzed pregnancies at risk of mutations related to hemophilia A, hemophilia B and ornithine transcarbamylase deficiency (OTC). We designed Taqman assays targeting these mutations (Supp. Table S3) and validated them as explained in the methods section. We then ran the validated assay for each sample and counted the Poisson corrected number of mutated (N_M_) and healthy (N_H_) alleles in maternal plasma (Fig. 4a-b). From the measured fetal fraction of each sample, we determined the ratio of mutated and healthy alleles that we would expect for an affected or an unaffected pregnancy as well as its associated error (Supp. Section S2.1) ^25^. We used this information to compute the expected distributions for an affected or an unaffected pregnancy and compare them to the experimentally measured ratio (Fig. 4g-h, blue and green distributions and dotted arrow). The affected or unaffected status of the fetus was determined from the probability of the measurement arising from each distribution using a likelihood ratio classifier (Fig 4g-h and Methods) ^27,31^. Using this approach, we also analyzed two pregnancies at-risk of OTC. First we tested a non-carrier mother at-risk due to gamete mosaicism (detected through a previously affected sibling), which we determined to be an unaffected pregnancy (Supp. Fig. S5). Then we analyzed a pregnancy carrying a female fetus that we determined to be a carrier of the maternal mutation, and therefore at a partial risk of post-neonatal-onset (Supp. Fig. S6).

**Figure 4.**
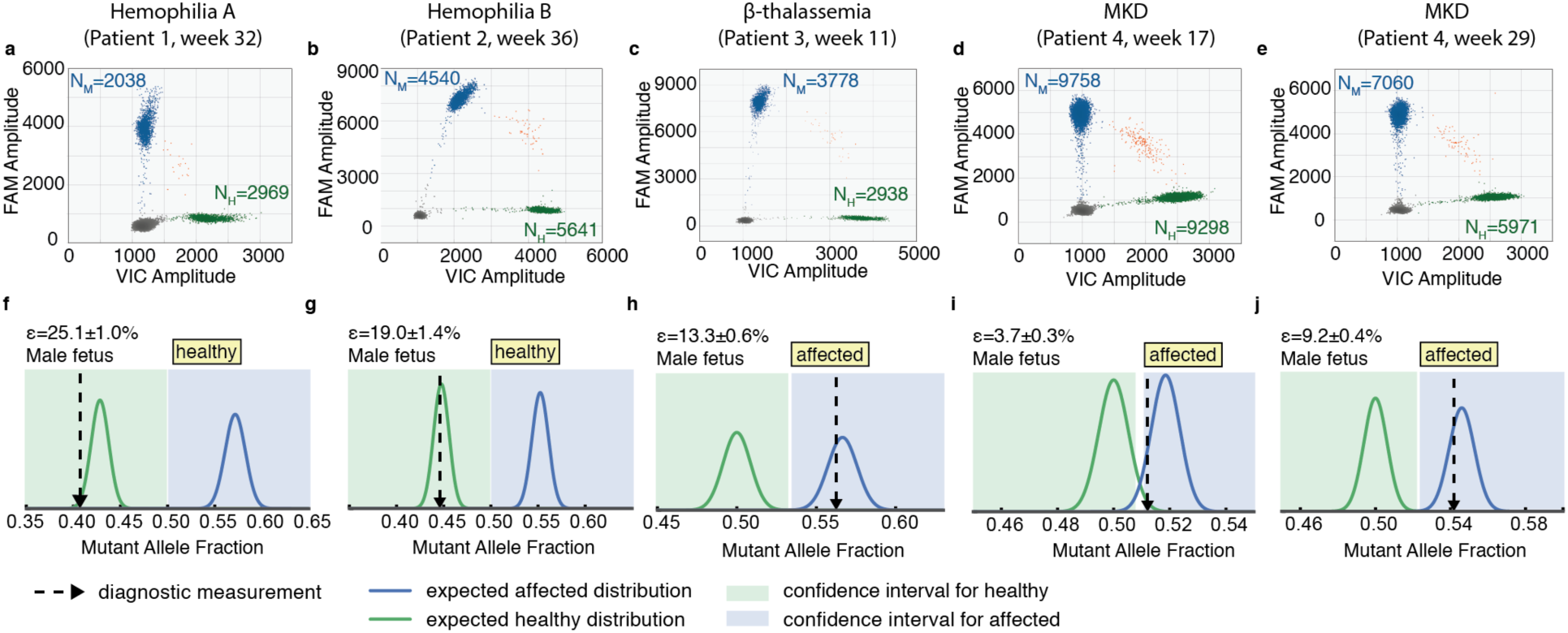
Diagnosis of fetuses at risk of maternally-inherited mutations (a-e) **(a)** Measurement of total counts of mutant (FAM) and healthy (VIC) alleles in maternal plasma using ddPCR for 5 different samples at risk of (a) Hemophilia A (b) Hemophilia B, (c) β-thalassemia and (d,e) mevalonate kinase deficiency. Clusters correspond to droplets positive for the mutant allele (blue), the healthy allele (green), both alleles (orange) or none (gray). N_M_ and N_H_ are the Poisson corrected counts for the mutant and healthy alleles respectively. **(f-j)** Diagnostic test for each sample a-e using a likelihood ratio classifier (see Methods). The dotted arrow corresponds to the measured ratio of mutant allele. The expected distributions for a sample with fetal fraction ε and carrying a healthy (affected) fetus is plotted in green (blue). The areas shaded in green and blue correspond to the ratios for which a fetus is determined to be healthy or affected using the ratio classifier (see Methods). Fetal fraction ε is reported as mean±SEM.

We then addressed the case of autosomal recessive mutations where both mother and father are carriers of the same mutation and therefore at a 25% risk of having an affected pregnancy. We analyzed pregnancies at risk of β-thalassemia and mevalonate kinase deficiency (MKD). To perform the assay, we followed the same approach described for X-linked mutations but used the counts and distributions expected for an autosomal recessive disorder (Supp. Section S2.2) ^24,26,30^. We ran the assay for each maternal plasma sample and measured the number of mutated and healthy alleles (Fig. 4c-d). For both samples the measured ratio was within the confidence intervals for an affected pregnancy (Fig. 4h-i). The affected status of the MKD case was also confirmed in a sample collected later in pregnancy and having a higher fetal fraction (Fig. 4e-j). All measurements were also confirmed in postnatal testing and found to be in agreement with the non-invasive prenatal test.

### Diagnosis of heterozygous compound mutations

Finally, we address the case of single gene disorders where each parent carries a different mutation affecting the same gene. We first tested pregnancies at risk of muscle-type acetylcholine receptor (AChR) deficiency (mutations: c459dupA and c753_754delAA), and cystic fibrosis (mutations: ΔF508 and W1282X). The latter are the two most common mutations for cystic fibrosis in Ashkenazi Jews, with an estimated combined abundance >75% ^32^. For these conditions we ran the assay for the paternal mutation using enough sample to observe ∼40 counts of the mutated allele in an affected pregnancy. To determine this value we used the combined information of the fetal fraction and total cfDNA abundance in maternal plasma. From Poisson statistics, this sets the expected result for a fetus that is a carrier of the mutation approximately 6 standard deviations away from a negative result (p<10^-12^, see Methods). The remaining sample was used to detect inheritance of the maternal mutation (Supp. Section S2.2). For each sample we measured N_M_ and N_H_ for each mutation at-risk (Fig. 5a-d), and determined the genotype of the fetus from the probability of each measurement arising from a carrier or non-carrier using a likelihood ratio classifier (Fig. 5e-h). Both pregnancies were determined to have an unaffected fetus, although the fetus at risk of AChR deficiency was determined to be carrier of the maternal mutation whereas the fetus at risk of cystic fibrosis was determined to be carrier of the paternal mutation. Using this approach, we also analyzed a pregnancy at risk of GJB-2 related DFNB1 nonsyndromic hearing loss (mutations: c.71G> A and c.-23+1G> A) at week 16 of gestation (fetal fraction: 6.7±0.5%), which we determined not to be a carrier of the mutated alleles (Supp. Fig. S7).

**Figure 5.**
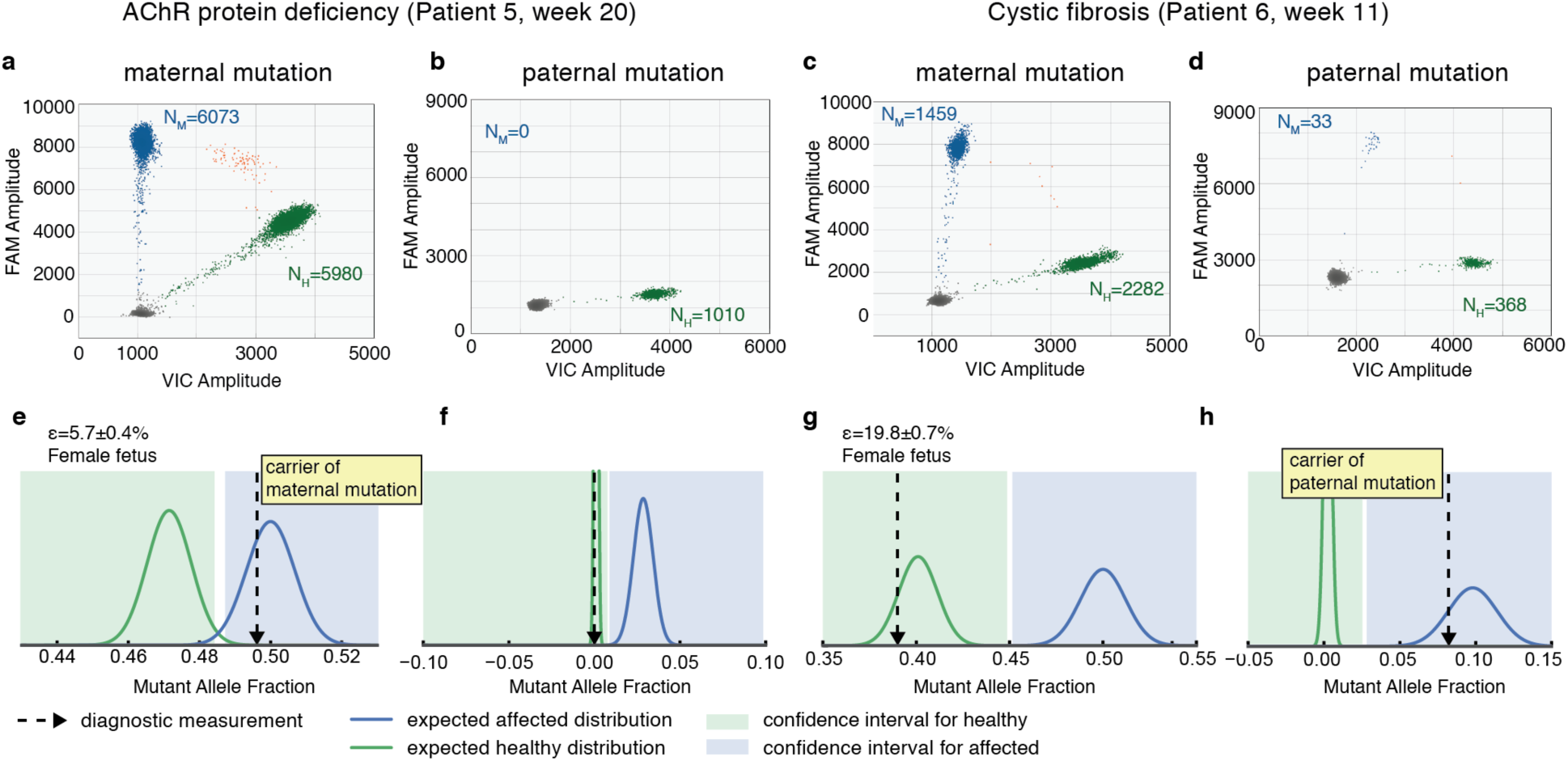
Diagnosis of fetuses at risk of combined paternal and maternal mutations for the same gene. **(a-d)** Measurement of total counts of mutant (FAM) and healthy (VIC) alleles in maternal plasma using ddPCR for a pregnancy at risk of (a,b) AchR deficiency and (c,d) cystic fibrosis. Panels (a) and (c) correspond to the assay testing inheritance of the maternal mutation; panels (b) and (d) correspond to the assay testing inheritance of the paternal mutation. Clusters correspond to droplets positive for the mutant allele (blue), the healthy allele (green), both alleles (orange) or none (gray). N_M_ and N_H_ are the Poisson corrected counts for the mutant and healthy alleles respectively. **(e-h)** Diagnostic test for the inheritance of each mutation in a-d using a likelihood ratio classifier (see Methods). The dotted arrow corresponds to the measured ratio of mutant allele. The expected distributions for a sample with fetal fraction ε and carrying a healthy (affected) fetus is plotted in green (blue). The areas shaded in green and blue correspond to the ratios for which a fetus is determined to be healthy or affected using a ratio classifier (see Methods). Fetal fraction ε is reported as mean±SEM.

## Discussion

In this work we developed a direct ddPCR approach to test pregnancies at risk of X-linked and autosomal recessive single-gene disorders both for single mutations and compound heterozygous mutations. The proposed protocol builds on previous work carried by other labs and our own ^25–28^, and does not require extensive sample preparation or computational resources, being significantly simpler and cost-effective than approaches using deep sequencing and maternal haplotyping ^29,33^. We show that using this approach, noninvasive prenatal diagnosis can be performed in a clinical laboratory setting in ∼1 day from sample collection. This is particularly relevant for single-gene disorders where samples typically come in sparsely and are rarely at-risk for the same mutation. We validated the approach by correctly diagnosing pregnancies at risk of some of the most common point mutations such as ΔF508 (accounting for >70% of cystic fibrosis cases in Europe) ^22,34^, as well as rare metabolic and neuromuscular disorders that had not yet been addressed using non-invasive techniques (e.g. OTC, MKD or AchR deficiency). Early diagnosis of these metabolic disorders would improve management of the disease, especially in cases where early onset might lead to the accumulation of metabolites that cause irreversible organ damage and failure ^35–37^.

The volume of blood collected for this study (30 ml.) is sufficient for accurate classification of samples with a fetal fraction down to 4%, which is comparable to current standards for NIPT ^38,39^. We have compiled a table of expected test performance as a function of fetal fraction and blood draw (Supp. Table S4), and these numbers are consistent with what we actually measure. For instance, in our most challenging sample (patient 4, fetal fraction 3.6%, Fig. 4d), we used the whole sample and obtained ∼20,000 counts, which is in agreement with our blood draw (∼25/30 ml blood) and within a range of expected type I and type II errors of 0.2-1%. For certain single gene disorders, the sensitivity of the assay could be increased by following high-variability SNPs close to the target mutation (using a similar multiplexing approach as the one used here for the fetal fraction determination). This might become a cost-effective approach for common conditions such as cystic fibrosis, although for many mutations the collection of a moderate volume of blood (2 Streck tubes), should enable a correct classification of samples down to a 5% fetal fraction with false positive and false negative rates ∼0.2% (in line with current standards for trisomy 21).

Finally, we also provide an accurate method to measure the fetal fraction and total amount of cfDNA in plasma samples using a multiplexed SNP panel for ddPCR. This approach is useful to establish confidence intervals in NIPT of autosomal recessive or X-linked diseases. As shown here, NIPT of these conditions relies on comparing the ratio of mutated and healthy alleles in maternal plasma to the ratios expected for a healthy or affected fetus as determined from the sample fetal fraction. Overall, the use of a SNP panel instead of a single marker to measure fetal fraction (i) reduces false positive and negative rates, (ii) reduces sample dropout due to a lack of indicative markers, and (iii) simplifies the workflow as an initial maternal genotyping step is not needed. Moreover, using our SNP panel we show that the errors associated with counting statistics at the mutation site and errors associated with fetal fraction determination are comparable in size, highlighting the importance of accurately measuring the sample fetal fraction to implement NIPT of single gene disorders.

## Materials and Methods

### Sample collection and cfDNA extraction

A total of 10 blood samples were collected from pregnancies at risk of a single-gene disorder. Samples were collected in cfDNA Streck tubes (3 tubes, approximately 30 mL). Blood was centrifuged at 1,600g for 10 minutes, and the supernatant was centrifuged for an additional 10 minutes at 16000g to remove cellular debris. Plasma samples were aliquoted in 2 ml tubes and stored at −80°**C** until further processing (cfDNA extraction). Maternal genomic DNA was extracted from the remaining cellular fraction using the Qiagen Blood Mini kit (200 μl aliquots), and stored for assay validation. Extraction of cfDNA from stored plasma samples was done using the Qiagen Circulating Nucleic Acid kit using the protocol recommended by the manufacturer with the following modifications: we performed an initial centrifugation of plasma for 3 minutes at 14,000 rpm to remove cryoprecipitates, we extended the lysis step to 1h (as recommended for Streck tubes), and we did not add carrier RNA as we did not observe an increase in yield. Plasma was processed in batches of 5 ml per Qiagen column and eluted in 50 μl TE buffer.

### Quantification of cfDNA in plasma and fetal fraction

From the extracted cfDNA (∼150 μl in total) we used 8.5 μl (-850 μl plasma) for a preamplification reaction targeting highly variable SNPs that we used to determine the fetal fraction of each sample. We selected a total of 47 biallelic SNPs that show a high minor allele fraction (MAF > 0.4) for all five superpopulations of the 1000 genomes project (EAS, EUR, AFR, AMR, SAS) and that are not found in regions of structural variation or highly repetitive regions (filtered using UCSC RepeatMasker and the Database of Genomic Variants). Commercially available SNP Genotyping assays (ThermoFisher) were purchased for the selected SNPs (amplicon size < 80 bp), as well as separate primers targeting each SNP region (Supp. Table S1). An additional SNP Taqman assay targeting the ZFX and ZFY genes in chromosomes X/Y was also included in the assay ^40^. The size of the SNP panel, threshold MAF, and chromosomal distribution of assays was designed to maximize the probability of making an accurate determination of the fetal fraction across a broad target population (Supp. Fig. S3).

The preamplification reaction was performed using the Taqman PreAmp Master Mix (Applied Biosystems, Ref. 4391128) with the pooled 48 primer pairs and the recommended conditions by the manufacturer (reaction volume 50 μl, final primer concentration 45 nM each, 11 preamplification cycles). The preamplified DNA was diluted 5X with TE buffer and stored for ddPCR quantification.

Quantification of the fetal fraction and total amount of cfDNA were performed using ddPCR and standard conditions (reaction volume 20μl, final primer (probe) concentration: 900 nM (200 nM), thermal cycling: [10’ 95°**C**; 40X[30” 94°**C**;1’ 60°**C**]; 10’ 98°**C**], ramp rate: 2°**C**/sec). We used 1 μl of the preamplified DNA for each SNP Taqman assay reaction and 1 μl of the original cfDNA for each quantification assay (Supp. Tables S1 and S2). For the quantification assays we used 2 multiplex assays targeting chromosomes 1, 5, 10 and 14 (Supp. Table S2). Fetal fraction and quantification ddPCR assays were run in parallel in a single plate.

The amount of cfDNA per ml of plasma (in genomic equivalents) is determined as the mean of the four quantification assays. For the SNP assays, Poisson corrected counts are determined as *N*_FAM/VIC_ = -*N*_total_ *ln*[1-*N*_positive_/*N*_total_] (Equation 1), where *N*_*total*_ is the total number of droplets and *N*_*positive*_ the number of positive droplets for each channel (FAM or VIC). For each SNP assay the minor allele fraction is extracted as MAF = min(*N*_FAM_,*N*_VIC_)/(*N*_FAM_ + *N*_VIC_). The fetal fraction (ε) is determined from the median of all SNPs where the fetus is heterozygous and the mother homozygous (0.5% > MAF > 20%) using. Errors are determined as the standard deviation (SD) and compared to the Poisson noise expected from the DNA input used in the preamplification reaction 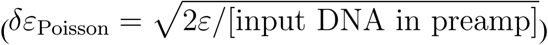.

### Results on Clinical Samples

For each sample we initially measured the fetal fraction and total amount of cfDNA as detailed above. From these values we determined the optimal split of sample between the paternal and maternal mutation, as well as the probability of obtaining an unambiguous result. Assays to detect inheritance of the mutations were designed and validated as described in Supp. Section S1. To test the paternal mutation we used the amount of cfDNA expected to provide ∼40 counts for a carrier fetus. This sets the result 6 standard deviations away from the non-carrier case. The remaining sample was used to quantify the imbalance on the maternal mutation. The ddPCR measurements were run using standard conditions and optimal temperatures determined in the validation assays. For each assay the total number of counts for each allele was determined using Equation 1. The affected or unaffected status of the fetus was determined using a likelihood ratio classifier with a low threshold of p(X|H_1_)/p(X|H_0_) = 1/8 and a high threshold of p(X|H_1_)/p(X|H_0_) = 8, where p(X|H_1_) is the probability of this result coming from an affected fetus and p(X|H_0_) is the null hypothesis of a non-affected fetus ^26,27,31^.

## Acknowledgments

We thank Ariana Spiegel, Anna Girsen and Katie Sherwin for research coordination, as well as Yasemin Dilara Sucu, Thuy Ngo and Kiran Kocherlota for help in collecting the plasma samples. This work was supported by the Hearst Foundation.

